# Robust and Structural Ergodicity Analysis and Antithetic Integral Control of a Class of Stochastic Reaction Networks

**DOI:** 10.1101/481051

**Authors:** Corentin Briat, Mustafa Khammash

## Abstract

Controlling stochastic reactions networks is a challenging problem with important implications in various fields such as systems and synthetic biology. Various regulation motifs have been discovered or posited over the recent years, a very recent one being the so-called Antithetic Integral Control (AIC) motif [3]. Several appealing properties for the AIC motif have been demonstrated for classes of reaction networks that satisfy certain irreducibility, ergodicity and output controllability conditions. Here we address the problem of verifying these conditions for large sets of reaction networks with time-invariant topologies, either from a robust or a structural viewpoint, using three different approaches. The first one adopts a robust viewpoint and relies on the notion of interval matrices. The second one adopts a structural viewpoint and is based on sign properties of matrices. The last one is a direct approach where the parameter dependence is exactly taken into account and can be used to obtain both robust and structural conditions. The obtained results lie in the same spirit as those obtained in [3] where properties of reaction networks are independently characterized in terms of control theoretic concepts, linear programs, and graph-theoretic/algebraic conditions. Alternatively, those conditions can be cast as convex optimization problems that can be checked efficiently using modern optimization methods. Several examples are given for illustration.

## 1 Introduction

The main objective of synthetic biology is the rational and systematic design of biological networks that can achieve de-novo functions such as the heterologous production of a metabolite of interest [4]. Besides the obvious necessity of developing experimental methodologies allowing for the reliable implementation of synthetic networks, tailored theoretical and computational tools for their design, simulation, analysis and control also need to developed. Indeed, theoretical tools that could predict certain properties (e.g. a stable/oscillatory/switching behavior, controllable trajectories, etc.) of a synthetic biological network from an associated model formulated, for instance, in terms of a reaction network [5–7], could pave the way to the development of reliable iterative procedures for the systematic design of efficient synthetic biological networks. Such an approach would allow for a faster design procedure than those involving fastidious experimental steps, and would give insights on how to adapt the current design in order to improve a certain design criterion. This way, synthetic biology would become conceptually much closer to existing theoretically-driven engineering disciplines, such as control engineering. However, while such methods are well-developed for deterministic models (i.e. deterministic reaction networks), they still lag behind in the stochastic setting. This lack of tools is quite problematic since it is now well-known that stochastic reaction networks [7] are versatile modeling tools that can capture the inherent stochastic behavior of living cells [8,9] and can exhibit several interesting properties that are absent for their deterministic counterparts [3, 10–12]. Under the well-mixed assumption, it is known [13, 14] that such random dynamics can be well represented as a continuous-time jump Markov process evolving on the *d*-dimensional nonnegative integer lattice where *d* is the number of distinct molecular species involved in the network. Sufficient conditions for checking the ergodicity of open unimolecular and bimolecular stochastic reaction networks has have been proposed in [15] and formulated in terms of linear programs. The concept of ergodicity is of fundamental importance as it can serve as a basis for the development of a control theory for biological systems. Indeed, verifying the ergodicity of a control system, consisting for instance of an endogenous biomolecular network controlled by a synthetic controller (see Fig. 1**A**), would prove that the closed-loop system is well-behaved (e.g. ergodic with bounded first- and second-order moments) and that the designed control system achieves its goal (e.g. set-point tracking and perfect adaptation properties). This procedure is analogous to that of checking the stability of a closed-loop system in the deterministic setting; see e.g. [16]. Additionally, designing synthetic circuits achieving a given function that are provably ergodic could allow for the rational design of synthetic networks that can exploit noise in their function. A recent example is that of the antithetic integral feedback controller proposed in [3] (see also Fig. 1**B**) that has been shown to induce an ergodic closed-loop network when some conditions on the endogenous network to be controlled are met; see also [17, 18].

**Figure 1:**
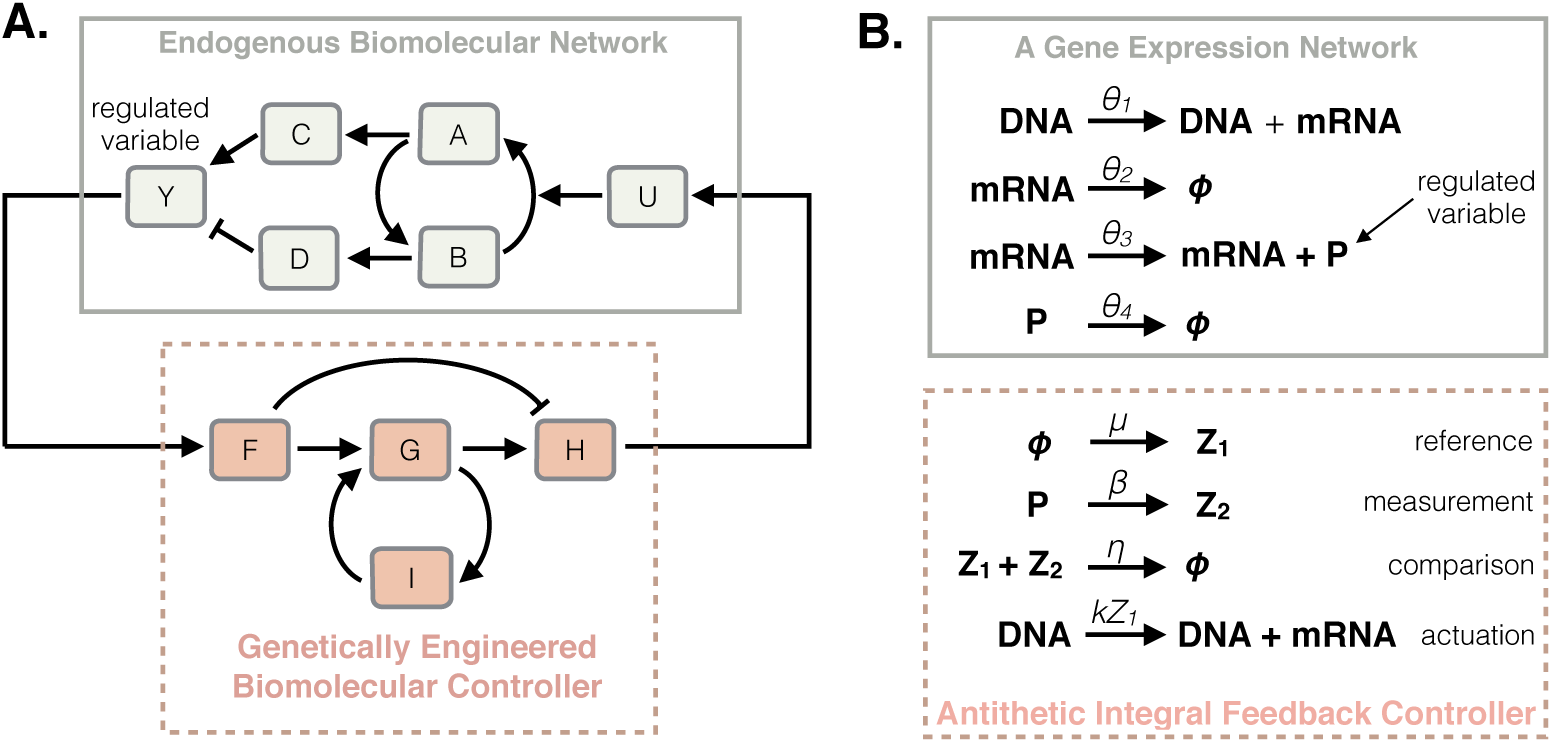
**A**. A synthetic feedback loop involving an endogenous network controlled with a synthetic feedback controller. **B**. A gene expression network (top) and an antithetic integral controller (bottom) as examples of endogenous and synthetic networks.

A major limitation of the ergodicity conditions obtained in [3, 15, 19] is that they only apply to networks with fixed and known rate parameters – an assumption that is rarely met in practice as the rate parameters are usually poorly known and context dependent. This motivates the consideration of networks with uncertain rate parameters [1, 2]. Three approaches are discussed in this paper. The first one is quantitative and considers networks having an interval matrix [20] as characteristic matrix. We show that checking the ergodicity and the output controllability of the entire network class reduces to checking the Hurwitz stability of a single matrix and the output controllability of a single positive linear system. We also show that these conditions exactly write as a simple linear program having the same complexity as the program associated with the nominal case; i.e. in the case of constant and fixed rate parameters. The second approach is qualitative and is based on the theory of sign-matrices [21–24] which has been extensively studied and considered for the qualitative analysis of dynamical systems. Sign-matrices have also been considered in the context of reaction networks, albeit much more sporadically; see e.g. [1, 23, 25–28]. In this case, again, the conditions obtained in [1, 27] can be stated as a very simple linear program that can be shown to be equivalent to some graph theoretical conditions. A different approach relying on reaction network theory [5, 29] is also proposed in [30].

However, these approaches can be very conservative when the entries of the characteristic matrix of the network are not independent – a situation that appears when conversion reactions are involved in the networks or when reaction rates are functions of some hyper-parameters. In order to solve this problem, the actual parameter dependence needs to be exactly captured and many approaches exist to attack this problem such as, to cite a few, *µ*-analysis [31,32], small-gain methods [32,33], eigenvalue and perturbation methods [34,35], Lyapunov methods [36–40], etc. Unfortunately, it is well-known that checking whether a general parameter-dependent matrix is Hurwitz stable for all the parameter values is NP-Hard. In this regard, it seems important to develop an approach that is tailored to our problem by exploiting its inherent properties. By exploiting the structure of the problem at hand, several conditions for the robust ergodicity of unimolecular and biomolecular networks are first obtained in terms of a sign switching property for the determinant of the upper-bound of the characteristic matrix of the network. This condition also alternatively formulates as the existence of a positive vector depending polynomially on the uncertain parameters and satisfying certain inequality conditions. The complexity of the problem is notably reduced by exploiting the Metzler structure^1^ of the matrices involved and through the use of various algebraic results such as the Perron-Frobenius theorem. The structural ergodicity of unimolecular networks is also considered and shown to reduce to the analysis of constant matrices when some certain realistic assumptions are met. It is notably shown in the examples that this latter result can be applied to bimolecular networks in some situations.

### Outline

We recall in Section 2 several definitions and results related to reaction networks and antithetic integral control. Section 3 is concerned with the extension of the results in [3] to characteristic interval-matrices whereas Section 4 addresses the same problem using a parametric approach. Section 5 is concerned with the structural case where only the sign pattern of the characteristic matrix is known whereas Section 6 is devoted to the same problem using a parametric approach. Examples are treated in Section 7.

### Notations

The standard basis for 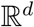 is denoted by 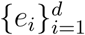. The sets of integers, non-negative integers, nonnegative real numbers and positive real numbers are denoted by 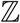, 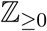, 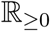 and 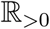, respectively. The *d*-dimensional vector of ones is denoted by 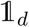 (the index will be dropped when the dimension is obvious). For vectors and matrices, the inequality signs ≤ and < act componentwise. Finally, the vector or matrix obtained by stacking the elements *x*_1_,…,*x_d_* is denoted by 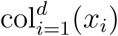 or col(*x*_1_,…,*x_d_*). The diagonal operator diag(·) is defined analogously. The spectral radius of a matrix 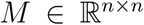 is defined as *ϱ*(*M*) = max{|*λ*| : det(*λI* − *M*) = 0}.

## 2 Preliminaries

### 2.1 Reaction networks

We consider here a reaction network with *d* molecular species ***X***_1_,…, ***X_d_*** that interacts through *K* reaction channels 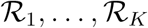 defined as

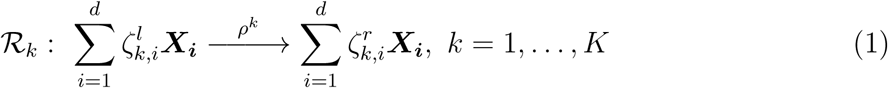

where 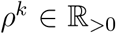 is the reaction rate parameter and 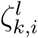, 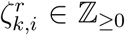. Each reaction is additionally described by a stoichiometric vector and a propensity function. The stoichiometric vector of reaction 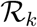 is given by 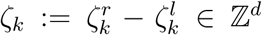 where 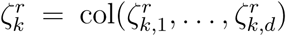 and 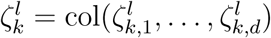. In this regard, when the reaction 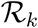 fires, the state jumps from *x* to *x* + *ζ_k_*. We define the stoichiometry matrix 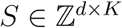 as *S* := [*ζ*_1_ … *ζ*_K_]. When the kinetics is mass-action, the propensity function of reaction 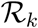 is given by 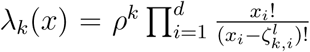 and is such that *λ_k_*(*x*) = 0 if 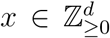 and 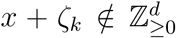. We denote this reaction network by 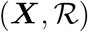. Under the well-mixed assumption, this network can be described by a continuous-time Markov process (***X***_1_(*t*),…,*X_d_*(*t*))*_t_*_≥0_ with state-space 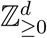; see e.g. [13].

### 2.2 Ergodicity of unimolecular and bimolecular reaction networks

Let us assume here that the network 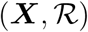 is at most bimolecular and that the reaction rates are all independent of each other. In such a case, the propensity functions are polynomials of at most degree 2 and we can write the propensity vector as

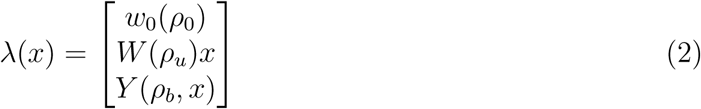

where 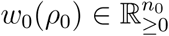, 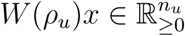 and 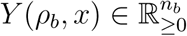 are the propensity vectors associated the zeroth-, first-and second-order reactions, respectively. Their respective rate parameters are also given by *ρ*_0_, *ρ_u_* and *ρ_b_*, and according to this structure, the stoichiometric matrix is decomposed as *S* =: [*S*_0_ *S_u_ S_b_*]. Before stating the main results of the section, we need to introduce the following terminology:

#### Definition 1

*The* characteristic matrix *A*(*ρ_u_*) *and the* offset vector *b*_0_(*ρ*) *of a bimolecular reaction network* 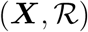 *are defined as*

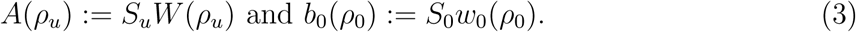

A particularity is that the matrix *A*(*ρ_u_*) is Metzler (i.e. all the off-diagonal elements are nonnegative) for all *ρ_u_* ≥ 0. This property plays an essential role in the derivation of the results of [3] and will also be essential for the derivation of the main results of this paper. It is also important to define the property of ergodicity:

#### Definition 2

*The Markov process associated with the reaction network* 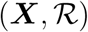 *is said to be ergodic if its probability distribution globally converges to unique stationary distribution. It is exponentially ergodic if the convergence to the stationary distribution is exponential*.

We then have the following result:

#### Theorem 3

**([15])** *Let us consider an irreducible*^2^ *bimolecular reaction network* 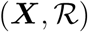 *with fixed rate parameters; i.e. A* = *A*(*ρ_u_*) *and b*_0_ = *b*_0_(*ρ*_0_)*. Assume that there exists a vector* 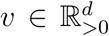 *such that v^T^ S_b_* = 0 *and v^T^ A* < 0*. Then, the reaction network* 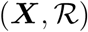 *is exponentially ergodic and all the moments are bounded and converging*.

We also have the following immediate corollary pertaining on unimolecular reaction networks:

#### Corollary 4

*Let us consider an irreducible unimolecular reaction network* 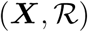 *with fixed rate parameters; i.e. A* = *A*(*ρ_u_*) *and b*_0_ = *b*_0_(*ρ*_0_)*. Assume that there exists a vector* 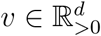 *such that v^T^ A<* 0*. Then, the reaction network* 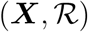 *is exponentially ergodic and all the moments are bounded and converging*.

### 2.3 Antithetic integral control of unimolecular networks

Antithetic integral control has been first proposed in [3] for solving the perfect adaptation problem in stochastic reaction networks. The underlying idea is to augment the open-loop network 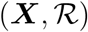 with an additional set of species and reactions (the controller). The usual set-up is that this controller network acts on the production rate of the molecular species ***X***_1_ (the *actuated species*) in order to steer the mean value of the *controlled species* ***X_ℓ_***, *ℓ* ∈ {1,…,*d*}, to a desired set-point (the reference). To the regulation problem, it is often sought to have a controller that can ensure perfect adaptation for the controlled species. As proved in [3], the antithetic integral control motif 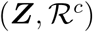 defined with

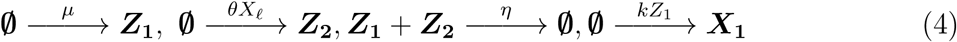

solves this control problem with the set-point being equal to *µ*/*θ*. Above, ***Z*_1_** and ***Z*_2_** are the *controller species*. The four controller parameters *µ,θ,η,k* > 0 are assumed to be freely assignable to any desired value. The first reaction is the *reference reaction* as it encodes part of the reference value *µ/θ* as its own rate. The second one is the *measurement reaction* that produces the species ***Z*_2_** at a rate proportional to the current population of the controlled species ***X_ℓ_***. The third reaction is the *comparison reaction* as it compares the populations of the controller species and annihilates one molecule of each when these populations are both positive. Finally, the fourth reaction is the *actuation reaction* that produces the actuated species ***X*_1_** at a rate proportional to the controller species ***Z*_1_**.

The following fundamental result states conditions under which a unimolecular reaction network can be controlled using an antithetic integral controller:

#### Theorem 5

**( [3])** *Suppose that the open-loop reaction network* 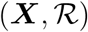 *is unimolecular and that the state-space of the closed-loop reaction network* 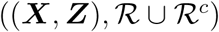 *is irreducible. Let us assume that ρ*_0_ *and ρ_u_ are fixed and known (i.e. A* = *A*(*ρ_u_*) *and b*_0_ = *b*_0_(*ρ*_0_)*) and assume, further, that there exist vectors* 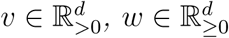*, w*_1_ > 0*, such that*

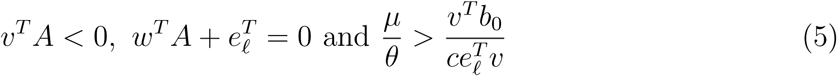

*where c>* 0 *verifies v^T^* (*A* + *cI*) ≤ 0.

*Then, for any values for the controller rate parameters η,k* > 0*, (i) the closed-loop network is ergodic, (ii)* 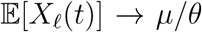 *as t* → ∞ *and (iii)* 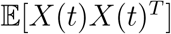 *is bounded over time*.

We can see that the conditions above consist of the combination of an ergodicity condition (*v^T^ A* < 0) and an output controllability condition for Hurwitz stable matrices *A* (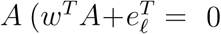 with *w*_1_ > 0), which are fully consistent with the considered control problem. Note, however, that unlike in the deterministic case, the above result proves that the closed-loop network cannot be unstable; i.e. have trajectories that grow unboundedly with time. This is illustrated in more details in the supplemental material of [3].

We will also need the following result on the output controllability of linear SISO positive systems:

#### Lemma 6

**( [3])** *Let* 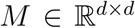 *be a Metzler matrix. Then, the following statements are equivalent:*

a. *The linear system*

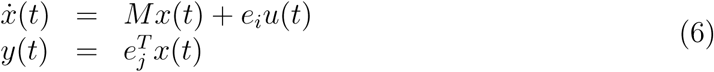

*is output controllable*.
b. rank 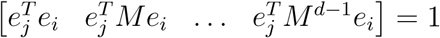.
c. *There is a path from node i to node j in the directed graph* 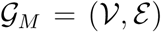 *defined with* 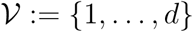 *and*

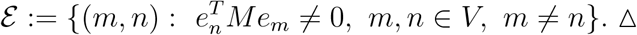 *Moreover, when the matrix M is Hurwitz stable, then the above statements are also equivalent to:*
d. *The inequality* 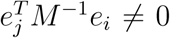 *holds or, equivalently, the static-gain of the system* (6) *is nonzero*.

Before stating the main result, it is convenient to define here the following properties:

**P1**. the closed-loop reaction network 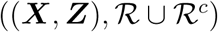 is ergodic,

**P2**. the mean of the controlled species satisfies 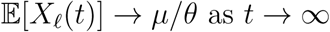 as *t* →∞,

**P3**. the second-order moment matrix 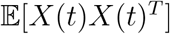 is uniformly bounded and globally converges to its unique stationary value.

We are now ready to state the main result of [3] on unimolecular networks:

#### Theorem 7

**( [3])** *Assume that the open-loop reaction network* 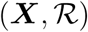 *is unimolecular and that the state-space of the closed-loop reaction network* 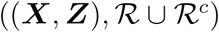 *is irreducible. Let us also assume that the vector of reaction rates ρ is fixed and equal to some nominal value* 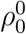 *and* 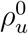 *for the zeroth- and first-order reactions, respectively. In this regard, we set* 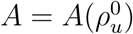 *and* 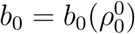*. Then, the following statements are equivalent:*

a. *There exist vectors* 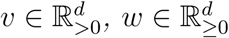*, w*_1_ > 0*, such that v^T^ A<* 0 *and* 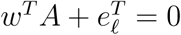.
b. *The positive linear system describing the dynamics of the first-order moments given by*

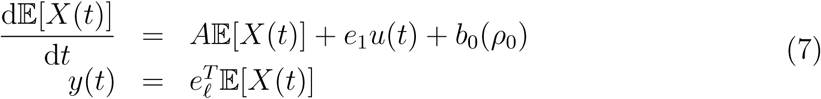

*is asymptotically stable and output controllable; i.e. the characteristic matrix A of the network* 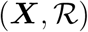 *is Hurwitz stable and*

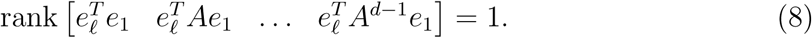 *Moreover, when one of the above statements holds, then for any values for the controller rate parameters η,k* > 0*, the properties* ***P1., P2***. *and* ***P3***. *hold provided that*

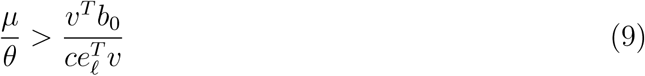

*where c>* 0 *and* 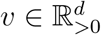 *verify v^T^* (*A* + *cI*) ≤ 0.

Interestingly, the conditions stated in the above result can be numerically verified by checking the feasibility of the following linear program

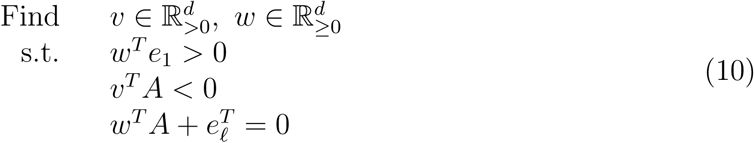

which involves 2*d* variables, 3*d* inequality constraints and *d* equality constraints.

## 3 Robust ergodicity of reaction networks - Interval matrices

### 3.1 Preliminaries

The results obtained in the previous section apply when the characteristic matrix 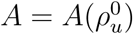 is fixed as the linear programming problem (10) can only be solved for that characteristic matrix. The goal is to generalize these results to the case where the characteristic matrix *A*(*ρ*) and the offset vector *b*_0_(*ρ*) are uncertain and belong to the sets

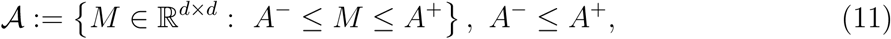

and

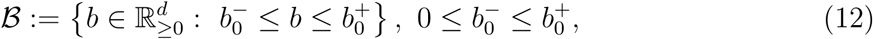

where the inequality signs are componentwise and the extremal Metzler matrices *A*^+^,*A*^−^ and vectors 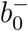, 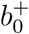 are known. These bounds can be determined such that the inequalities

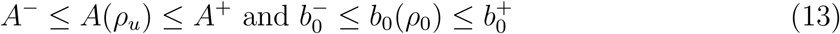

hold for all 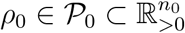 and 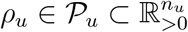 where 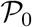 and 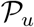 are compact sets containing all the possible values for the rate parameters. We have the following preliminary result:

#### Lemma 8

*The following statements are equivalent:*

a. *All the matrices in* 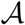 *are Hurwitz stable;*
b. *The matrix A*^+^ *is Hurwitz stable*.

#### Proof

The proof that (a) implies (b) is immediate. The converse can be proved using the fact that for two Metzler matrices *M*_1_, 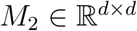 verifying the inequality *M*_1_ ≤ *M*_2_, we have that *λ_F_* (*M*_1_) ≤ *λ_F_* (*M*_2_) where *λ_F_* (·) denotes the Frobenius eigenvalue (see e.g. [42]). Hence, we have that *λ_F_* (*M*) ≤ *λ_F_* (*A*^+^) < 0 for all 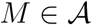. The conclusion then readily follows.

### 3.2 Main result

We are now in position to state the following generalization of Theorem 7:

#### Theorem 9

*Let us consider a unimolecular reaction network* 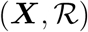 *with characteristic matrix A in* 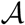 *and o*ff*set vector b*_0_ *in* 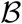*. Assume also that the state-space of the closed-loop reaction network* 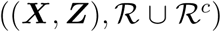 *is irreducible. Then, the following statements are equivalent:*

a. *All the matrices in* 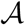 *are Hurwitz stable and for all* 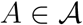*, there exists a vector* 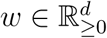 *such that w*_1_ > 0 *and* 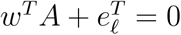.
b. *There exist two vectors* 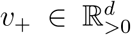, 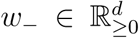 *such that* 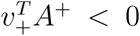, 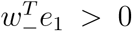 *and* 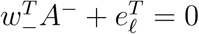. *Moreover, when one of the above statements holds, then for any values for the controller rate parameters η,k* > 0 *and any* 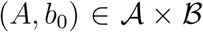*, the properties* ***P1., P2***. *and* ***P3***. *hold provided that*

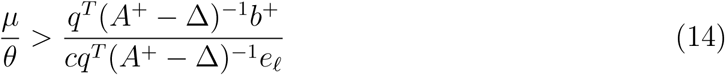

*and*

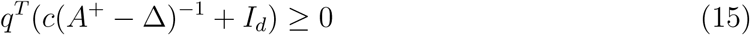

*for some c >* 0, 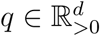 *and for all* Δ ∈ [0*,A*^+^ − *A*^−^].

#### Proof

The proof that (a) implies (b) is immediate. So let us focus on the reverse implication. Define, with some slight abuse of notation, the matrix *A*(Δ) := *A*^+^ − Δ, Δ ∈ **Δ**^+^ := [0*,A*^+^ − *A*^−^], where the set membership symbol is componentwise. The Hurwitz-stability of all the matrices in 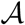 directly follows for the theory of linear positive systems and Lemma 8. We need now to construct a suitable positive vector 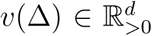 such that *v*(Δ)*^T^ A*(Δ) < 0 for all Δ ∈ **Δ^+^** provided that 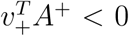. We prove now that such a *v*(Δ) is given by *v*(Δ) = (*I_d_* + Δ(*A*^+^ − Δ)^−1^)*^T^ v*_+_. Since *A*(Δ) = *A*^+^ − Δ, then we immediately get that 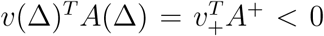. Hence, it remains to prove the positivity of the vector *v*(Δ) for all Δ ∈ **Δ**^+^. The difficulty here is that the product Δ(*A*^+^ − Δ)^−1^ is a nonnegative matrix since Δ ≥ 0 and (*A*^+^ − Δ)^−1^ ≤ 0, the latter being the consequence of the fact that *A*^+^ − Δ is Metzler and Hurwitz stable (see e.g. [42]). Therefore, there may exist values for 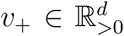 for which we have *v*(Δ) ≯ 0. To rule out this possibility, we restrict the analysis to all those 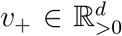 for which we have 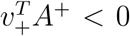. We can parameterize all these *v*_+_ as *v*_+_(*q*)= −(*A*^+^)^−^*^T^ q* where 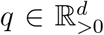 is arbitrary. We prove now that the vector *v*(Δ) = −(*I_d_* + Δ(*A*^+^ − Δ)^−1^)*^T^* (*A*^+^)^−^*^T^ q* > 0 is positive for all 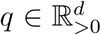 and for all Δ ∈ **Δ**^+^. This is is done by showing below that the matrix 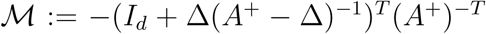 is nonnegative and invertible. Indeed, we have that

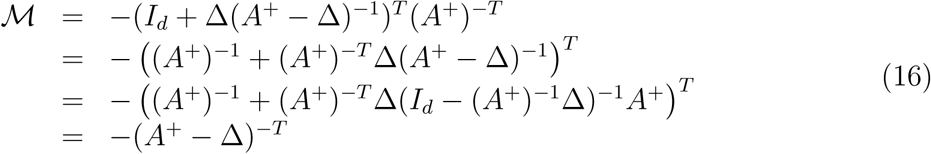

where the latter expression follows from the Woodbury matrix identity. Since (*A*^+^ − Δ) = *A*(Δ) is Metzler and Hurwitz stable for all Δ ∈ **Δ**^+^, then *A*^+^ − Δ is invertible and we have −(*A*^+^ − Δ)^−1^ ≥ 0, which proves the result.

Let us now consider then the output controllability condition and define *A*(Δ) as *A*(Δ) := *A*^−^ + Δ where Δ ∈ **Δ**^−^ := [0*,A*^+^ − *A*^−^]. We use a similar approach as previously and we build a *w*(Δ) that verifies the expression 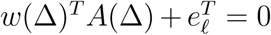 for all Δ ∈ **Δ**^−^ provided that 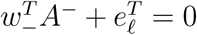. We prove that such a *w*(Δ) is given by *w*(Δ) := (*A*^−^(*A*^−^ + Δ)^−1^)*^T^ w*_−_. We first prove that this *w*(Δ) is nonnegative and that it verifies 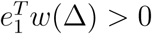 for all Δ ∈ **Δ**^−^. To show this, we rewrite this *w*(Δ) as *w*(Δ) = (*I_d_* − Δ(*A*^−^ + Δ)^−1^)*^T^ w*_−_ and using the fact that (*A*^−^ + Δ)^−1^ ≤ 0 since (*A*^−^ + Δ) is a Hurwitz stable Metzler matrix for all Δ ∈ **Δ**^−^, then we can conclude that *w*(Δ) ≥ *w*_−_ ≥ 0 for all Δ ∈ **Δ**^−^. An immediate consequence is that 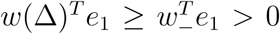 for all Δ ∈ **Δ**^−^. This proves the first part. We now show that this *w*(Δ) verifies the output controllability condition. Evaluating then *w*(Δ)*^T^* (*A*^−^ + Δ) yields

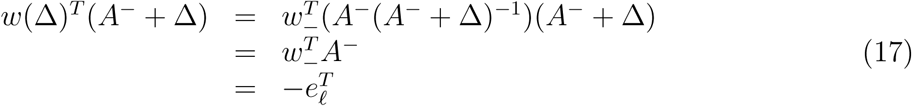

where the last row has been obtained from the assumption that 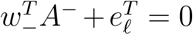. This proves the second part. Finally, the condition (14) is obtained by substituting the expression for *v*(Δ) defined above in (9). This completes the proof.

As in the nominal case, the above result can be exactly formulated as the linear program

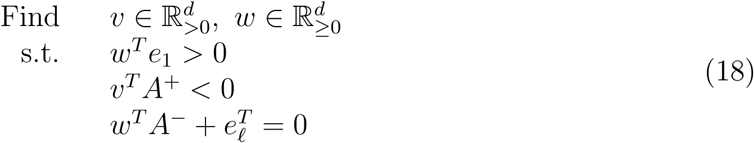

which has exactly the same complexity as the linear program (10). Hence, checking the possibility of controlling a family of networks defined by a characteristic interval-matrix is not more complicated that checking the possibility of controlling a single network.

### 3.3 A robust ergodicity result for bimolecular networks

We now provide an extension of the conditions of Theorem 3 for bimolecular networks to the case of uncertain networks described by uncertain matrices:

#### Proposition 10

*Let us consider the reaction network* 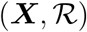 *and assume it is mass-action with at most reaction of order two. Assume further that the state-space of the underlying Markov process is irreducible and that there exists a vector* 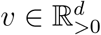 *such that*

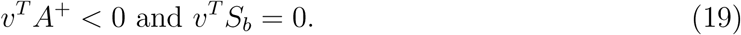

*Then, the stochastic reaction network* 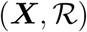 *is robustly exponentially ergodic for all A* ∈ [*A*^−^*,A*^+^].

#### Proof

The result immediately follows from Theorem 3 and Theorem 9.

Once again, the conditions can be efficiently checked by solving a linear program.

## 4 Robust ergodicity of reaction networks - Parametric approach

The approach based on interval matrices has the advantage of being very simple at the expense of some potentially high conservatism when the upper-bound *A*^+^ is not tight. This is the case when conversion reactions are involved. This may also be the case when some entries in the characteristic matrix are not independent; e.g. reaction rates are functions of some hyper-parameters. In this regard, the ergodicity test based on interval matrices may fail even if the characteristic matrix is Hurwitz stable for all possible values for the reaction rates. This motivates the consideration of a parametric approach tackling the problem in its primal form. Note that since the output observability test based on interval matrices is non-conservative, we only address here the problem of accurately checking the Hurwitz stability of the characteristic matrix for all admissible values for the reaction rates.

### 4.1 Preliminaries

The following lemma will be useful in proving the main results of this section:

#### Lemma 11

*Let us consider a matrix* 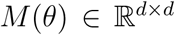 *which is Metzler and bounded for all* 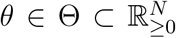 *and where* Θ *is assumed to be compact and connected. Then, the following statements are equivalent:*

a. *The matrix M*(*θ*) *is Hurwitz stable for all θ* ∈ Θ.
b. *The coefficients of the characteristic polynomial of M*(*θ*) *are positive* Θ.
c. *The conditions hold:*

(c1) *there exists a θ*^*^ ∈ Θ *such that M*(*θ*^*^) *is Hurwitz stable, and*
(c2) *for all θ* ∈ Θ *we have that* (−1)*^d^* det(*M*(*θ*)) > 0.

#### Proof

The proof of the equivalence between (a) and (b) follows, for instance, from [43] and is omitted. It is also immediate to prove that (b) implies (c) since if *M*(*θ*) is Hurwitz stable for all *θ* ∈ Θ then (c1) holds and the constant term of the characteristic polynomial of *M*(*θ*) is positive on *θ* ∈ Θ. Using now the fact that that constant term is equal to (−1)*^d^* det(*M*(*θ*)) yields the result.

To prove that (c) implies (a), we use the contraposition. Hence, let us assume that there exists at least a *θ_u_* ∈ Θ for which the matrix *M*(*θ_u_*) is not Hurwitz stable. If such a *θ_u_* can be arbitrarily chosen in Θ, then this implies the negation of statement (c1) (i.e. for all *θ*^*^ ∈ Θ the matrix *M*(*θ*^*^) is not Hurwitz stable) and the first part of the implication is proved.

Let us consider now the case where there exists some *θ_s_* ∈ Θ such that *M*(*θ_s_*) is Hurwitz stable. Let us then choose a *θ_u_* and a *θ_s_* such that *M*(*θ_u_*) is not Hurwitz stable and *M*(*θ_s_*) is. Since Θ is connected, then there exists a path 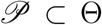 from *θ_s_* and *θ_u_*. From Perron-Frobenius theorem, the dominant eigenvalue, denoted by *λ_PF_*(·), is real and hence, we have that *λ_PF_*(*M*(*θ_s_*)) < 0 and *λ_PF_*(*M*(*θ_u_*)) ≥ 0. Hence, from the continuity of eigenvalues then there exists a 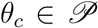 such that *λ_PF_*(*M*(*θ_c_*)) = 0, which then implies that (−1)*^d^* det(*M*(*θ_c_*)) = 0, or equivalently, that the negation of (c2) holds. This concludes the proof.

Before stating the next main result of this section, let us assume that *S_u_* in Definition 1 has the following form

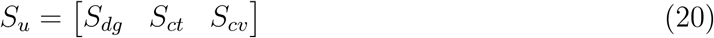

where 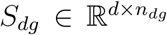 is a matrix with nonpositive columns, 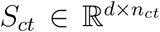 is a matrix with nonnegative columns and 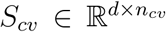 is a matrix with columns containing at least one negative and one positive entry. Also, decompose accordingly *ρ_u_* as *ρ_u_* =: col(*ρ_dg_, ρ_ct_, ρ_cv_*} and define

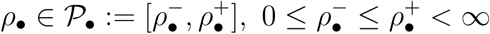

where •∈{*dg, ct, cv*} and let 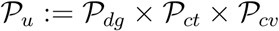.

In this regard, we can alternatively rewrite the matrix *A*(*ρ_u_*) as *A*(*ρ_dg_,ρ_ct_,ρ_cv_*). We then have the following result:

#### Lemma 12

*The following statements are equivalent:*

a. *The matrix A*(*ρ_u_*) *is Hurwitz stable for all* 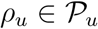.
b. *The matrix*

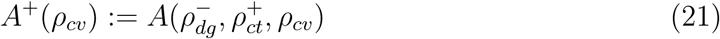

*is Hurwitz stable for all* 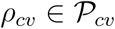.

#### Proof

The proof that (a) implies (b) is immediate. To prove that (b) implies (a), first note that we have

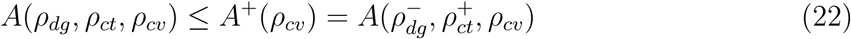

since for all 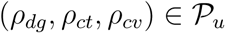. Using the fact that for two Metzler matrices *B*_1_, *B*_2_, the inequality *B*_1_ ≤ *B*_2_ implies *λ_PF_* (*B*_1_) ≤ *λ_PF_* (*B*_2_) [42], then we can conclude that 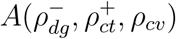 is Hurwitz stable for all 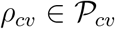 if and only if the matrix *A*(*ρ_dg_,ρ_ct_,ρ_cv_*) is Hurwitz stable for all 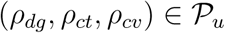. This completes the proof.

### 4.2 Unimolecular networks

The following theorem states the main result on the robust ergodicity of unimolecular reaction networks:

#### Theorem 13

*Let* 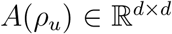 *be the characteristic matrix of some unimolecular network and* 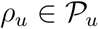*. Then, the following statements are equivalent:*

a. *The matrix A*(*ρ_u_*) *is Hurwitz stable for all* 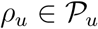.
b. *The matrix*

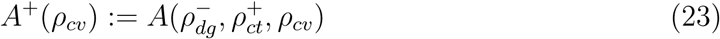

*is Hurwitz stable for all* 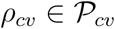.
c. *There exists a* 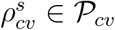 *such that the matrix* 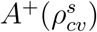 *is Hurwitz stable and the polynomial* (−1)*^d^* det(*A*^+^(*ρ_cv_*)) *is positive for all* 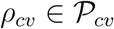.
d. *There exists a polynomial vector-valued function* 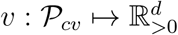 *of degree at most d* − 1 *such that v*(*ρ_cv_*)*^T^ A*^+^(*ρ_cv_*) < 0 *for all* 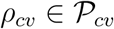.

#### Proof

The equivalence between the statement (a), (b) and (c) directly follows from Lemma 11 and Lemma 12. To prove the equivalence between the statements (b) and (d), first remark that (b) is equivalent to the fact that for any *q*(*ρ_cv_*) > 0 on 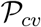, there exists a unique parameterized vector 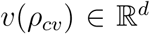 such that *v*(*ρ_cv_*) > 0 and *v*(*ρ_cv_*)*^T^* A^+^(*ρ_cv_*)= −*q*(*ρ_cv_*)*^T^* for all 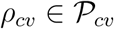. Choosing 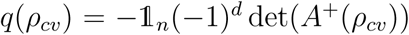, we get that such a *v*(*ρ_cv_*) is given by

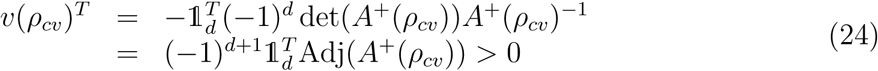

for all 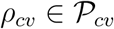. Since the matrix *A*^+^(*ρ_cv_*) is affine in *ρ_cv_*, then the adjugate matrix Adj(*A*^+^(*ρ_cv_*) contains entries of at most degree *d* − 1 and the conclusion follows.

Checking the condition (c) amounts to solving two problems. The first one is is concerned with the construction of a stabilizer 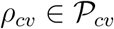 for the matrix *A*^+^(*ρ_cv_*) whereas the second one is about checking whether a polynomial is positive on a compact set. The first problem can be easily solved by checking whether *A*^+^(*ρ_cv_*) is Hurwitz stable for some randomly chosen point in 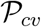. For the second one, optimization-based methods can be used such as those based on the Handelman’s Theorem combined with linear programming [44,45] or Putinar’s Positivstellensatz combined with semidefinite programming [46,47]. Note also that the degree *d* − 1 is a worst case degree and that, in fact, polynomials of lower degree will in general be enough for proving the Hurwitz stability of the matrix *A*^+^(*ρ_cv_*) for all 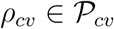. For instance, the matrices *A*(*ρ*) and *A*^+^(*ρ_cv_*) are very sparse in general due to the particular structure of biochemical reaction networks. The sparsity property is not considered here but could be exploited to refine the necessary degree for the polynomial vector *v*(*ρ_cv_*).

In is important to stress here that Theorem 13 can only be considered when the rate parameters are time-invariant (i.e. constant deterministic or drawn from a distribution). When they are time-varying (e.g. time-varying stationary random variables), a possible workaround relies on the use of a constant vector *v* as formulated below:

#### Proposition 14

(Constant *v*)

*Let* 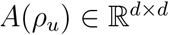 *be the characteristic matrix of some unimolecular network and* 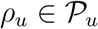*. Then, the following statements are equivalent:*

a. *There exists a vector* 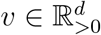 *such that v^T^ A*^+^(*ρ_cv_*) < 0 *holds for all* 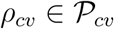.
b. *There exists a vector* 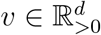 *such that v^T^ A*^+^(*θ*) < 0 *holds for all* 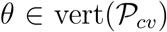 *where* 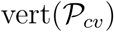 *denotes the set of vertices of the set* 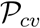.

#### Proof

The proof exploits the affine, hence convex, structure of the matrix *A*^+^(*ρ_cv_*). Using this property, it is indeed immediate to show that the inequality *v^T^ A*^+^(*ρ_cv_*) < 0 holds for all 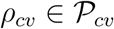 if and only if *v^T^ A*^+^(*θ*) < 0 holds for all 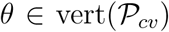 (see e.g. [36] for a similar arguments in the context of quadratic Lyapunov functions).

The above result is connected to the existence of a linear copositive Lyapunov function for a linear positive switched system with matrices in the family 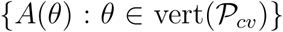 for which many characterizations exist; see e.g. [48, 49].

### 4.3 Bimolecular networks

In the case of bimolecular networks, we have the following result:

#### Proposition 15

*Let* 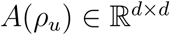 *be the characteristic matrix of some bimolecular network and* 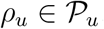*. Then, the following statements are equivalent:*

a. *There exists a polynomial vector-valued function* 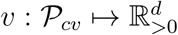 *of degree at most d* − 1 *such that*

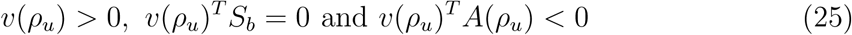

*for all* 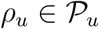.
b. *There exists a polynomial vector-valued function* 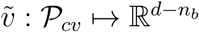 *of degree at most d* − 1 *such that*

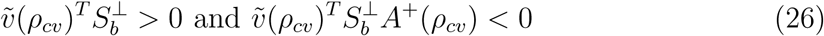

*for all* 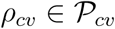 *and where n_b_* := rank(*S_b_*) *and* 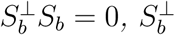 *full-rank*.

#### Proof

It is immediate to see that (a) implies (b). To prove the converse, first note that we have that 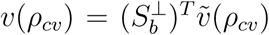 verifies *v*(*ρ_cv_*)*^T^ S_b_* = 0 and *v*(*ρ_cv_*) > 0 for all 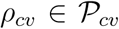. This proves the equality and the first inequality in (25). Observe now that for any 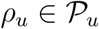 there exists a nonnegative matrix 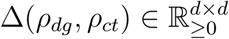 such that *A*(*ρ_u_*)= *A*^+^(*ρ_cv_*)−Δ(*ρ_dg_,ρ_ct_*). Hence, we have that

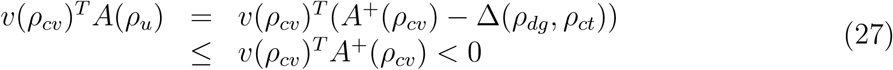

which proves the result.

As in the unimolecular case, we have been able to reduce the number of parameters by using an upper-bound on the characteristic matrix. It is also interesting to note that the condition 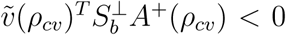 can be sometimes brought back to a problem of the form 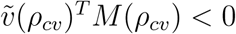 for some square, and often Metzler, matrix *M*(*ρ_cv_*) which can be dealt in the same way as in the unimolecular case.

The following result can be used when the parameters are time-varying and is the bimolecular analogue of Proposition 14:

#### Proposition 16 (Constant *v*)

*Let* 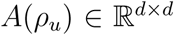 *be the characteristic matrix of some bimolecular network and* 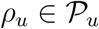*. Then, the following statements are equivalent:*

a. *There exists a vector* 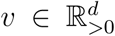 *such that v^T^ S_b_* = 0 *and v^T^ A*^+^(*ρ_cv_*) < 0 *hold for all* 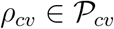.
b. *There exists a vector* 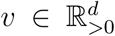 *such that v^T^ S_b_* = 0 *and v^T^ A*^+^(*θ*) < 0 *hold for all* 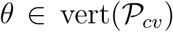.

## 5 Structural ergodicity of reaction networks network - Sign matrices

In the previous section, we were interested in uncertain networks characterized in terms of a characteristic interval-matrix. We consider here a different approach based on the qualitative analysis of matrices where we simply assume that we only know the sign-pattern of the characteristic matrix. We are then looking for criteria assessing whether all the characteristic matrices sharing the same sign-pattern verify the conditions of Theorem 7.

To this aim, let us consider the set of *sign symbols* 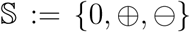 and define a *sign-matrix* as a matrix with entries in *S*. The qualitative class 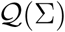 of a sign-matrix 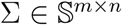 is defined as

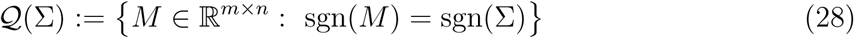

where the signum function sgn(·) is defined as

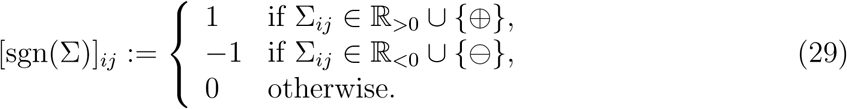

The following result proved in [27] will turn out to be a key ingredient for deriving the main result of this section:

#### Lemma 17

**( [27])** *Let* 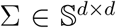 *be a given Metzler sign-matrix. Then, the following statements are equivalent:*

a. *All the matrices in* 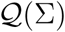 *are Hurwitz stable*.
b. *The matrix* sgn(Σ) *is Hurwitz stable*.
c. *The diagonal elements of* Σ *are negative and the directed graph* 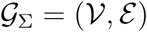 *defined with*

- 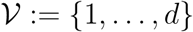 *and*
- 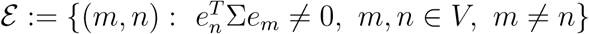.
d. *is an acyclic directed graph*.

We are now ready to state the main result of this section:

#### Theorem 18

*Let* 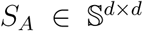 *be Metzler and S_b_* ∈ {0, ⊕}*^d^ be some given sign patterns for the characteristic matrix and the offset vector of some unimolecular reaction network* 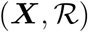*. Assume that ℓ* ≠ 1 *and that the state-space of the closed-loop reaction network* 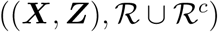 *is irreducible. Then, the following statements are equivalent:*

a. *All the matrices in* 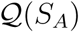 *are Hurwitz stable and, for all* 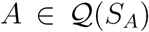*, there exists a vector* 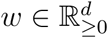 *such that w*_1_ > 0 *and* 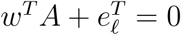.
b. *The diagonal elements of S_A_ are negative and the directed graph* 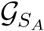 *is acyclic and contains a path from node* 1 *to node ℓ*.
c. *There exist vectors* 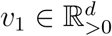, *v*_2_, 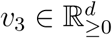, 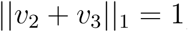*, such that the conditions*

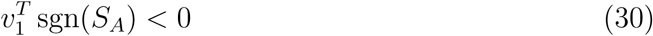

*and*

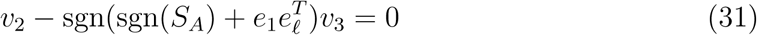

*hold*.

*Moreover, when one of the above statements holds, then for any values for the controller rate parameters η,k* > 0 *and any* 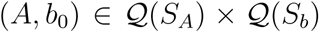*, the properties* ***P1., P2***. *and* ***P3***. *hold provided that*

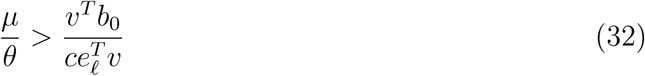

*where c>* 0 *and* 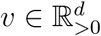 *verify v^T^* (*A* + *cI*) ≤ 0.

#### Proof

The equivalence between the statement (a) and statement (b) follows from Lemma 6 and Lemma 17. Hence, we simply have to prove the equivalence between statement (c) and statement (b).

The equivalence between the Hurwitz-stability of sgn(*S_A_*) and all the matrices in 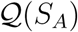 directly follows from Lemma 17. Note that if 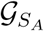 has a cycle, then there exists an unstable matrix in 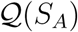. Let us assume then for the rest of the proof that there is no cycle in 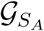 and let us focus now on the statement equivalent to the output controllability condition. From Lemma 17, we know that since all the matrices in 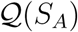 are Hurwitz stable, then the graph 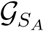 is an acyclic directed graph and *S_A_* has negative diagonal entries. From Lemma 6, we know that the network is output controllable if and only if there is a path from node 1 to node *ℓ* in the graph 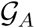. To algebraically formulate this, we introduce the sign matrix 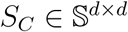 for which the associated graph 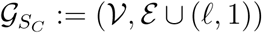, where *ℓ* ≠ 1 by assumption, consists of the original graph to which we add an edge from node *ℓ* to node 1. Note that if 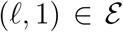, then *S_A_* = *S_C_*. The output controllability condition then equivalently turns into the existence of a cycle in the graph 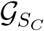 (recall the no cycle assumption for 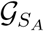 as, otherwise, some matrices in 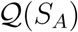 would not be Hurwitz stable). Considering again Lemma 6, we can turn the existence condition of a cycle in 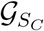 into an instability condition for some of the matrices in 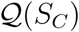. Since *S_C_* is a Metzler sign-matrix, then there exist some unstable matrices in 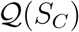 if and only if *v^T^* sgn(*S_C_*) ≮ 0 for all *v* > 0. Using Farkas’ lemma [50], this is equivalent to saying that there exist 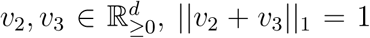 such that *v*_2_ − sgn(*S_C_*)*v*_3_ = 0. Therefore, the existence of *v*_2_*,v*_3_ verifying the conditions above is equivalent to saying that for all 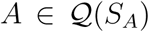, there exists a *w* ≥ 0, *w*_1_ > 0, such that 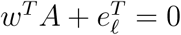. Noting, finally, that 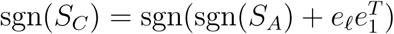 yields the result.

#### Remark 19

*In the case ℓ* = 1*, the output controllability condition is trivially satisfied as the actuated species coincides with the controlled species and hence we only need to check the Hurwitz stability condition v^T^* sgn(*S_A_*) < 0 *for some* 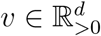.

As in the nominal and robust cases, the above result can also be naturally reformulated as the linear feasibility problem:

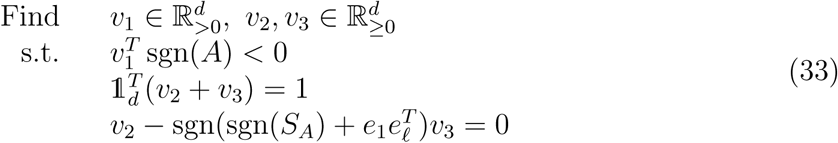

where 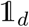 the *d*-dimensional vector of ones. The computational complexity of this program is slightly higher (i.e. 3*d* variables, 4*d* inequality constraints and 2*d* + 1 equality constraints) but is still linear in *d* and, therefore, this program will remain tractable even when *d* is large.

## 6 Structural ergodicity of reaction networks - Parametric approach

We are interesting in this section in the structural stability of the characteristic matrix of given unimolecular network. Hence, we have in this case 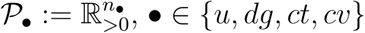 where *n*_•_ is the dimension of the vector *ρ*_•_.

### 6.1 A preliminary result

#### Lemma 20

*Let* 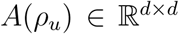 *be the characteristic matrix of some unimolecular network and* 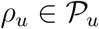*. Then, the following statements are equivalent:*

i. *For all* 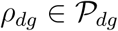 *and a* 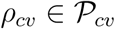*, the matrix A*(*ρ_dg_,ρ_cv_*, 0) *is Hurwitz stable*.
ii. *The matrix* 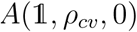 *is Hurwitz stable for all* 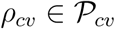.

#### Proof

The proof that (a) implies (b) is immediate. To prove the reverse implication, we use contraposition and we assume that there exist a 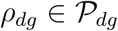 and a 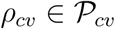 such that *A*(*ρ_dg_,ρ_cv_*, 0) is not Hurwitz stable. Then, we clearly have that

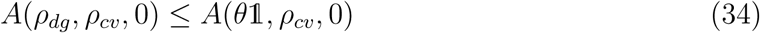

where *θ* = min(*ρ_dg_*) and hence 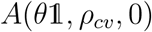 is not Hurwitz stable. Since 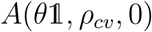 is affine in *θ* and *ρ_cv_*, then we have that 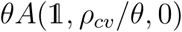 and since *θ* is independent of *ρ_cv_*, then we get that the matrix 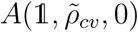 is not Hurwitz stable for some 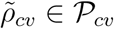. The proof is complete.

### 6.2 Main result

#### Theorem 21

*Let* 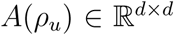 *be the characteristic matrix of some unimolecular network and* 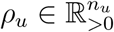*. Then, the following statements are equivalent:*

a. *he matrix A*(*ρ_u_*) *is Hurwitz stable for all* 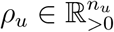.
b. *There exists a polynomial vector* 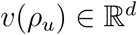 *of degree at most d* − 1 *such that v*(*ρ_u_*) > 0 *and v^T^ A*(*ρ_u_*) < 0 *for all* 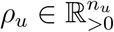.
c. *There exists a* 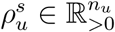 *such that the matrix* 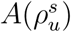 *is Hurwitz stable and the polynomial* (−1)*^d^* det(*A*(*ρ_u_*)) *is positive for all* 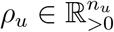.
d. *For all* 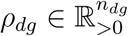 *and a* 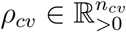*, the matrix A_ρ_* := *A*(*ρ_dg_,ρ_cv_*, 0) *is Hurwitz stable and we have that* 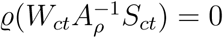.
e. *The matrix* 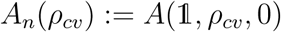 *is Hurwitz stable for all* 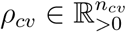 *and* 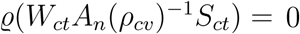 *for all* 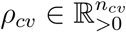. *Moreover, when S_cv_ contains exactly one entry equal to* −1 *and one equal to 1, then the above statements are also equivalent to*
f. *The matrix* 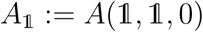 *is Hurwitz stable and* 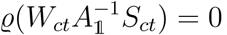.

#### Proof

The equivalence between the three first statements has been proved in Theorem 13. Let us prove now that (c) implies (d). Assuming that (c) holds, we get that the existence of a 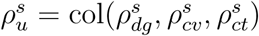 such that the matrix 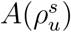 is Hurwitz stable immediately implies that the matrix *A_ρ_* = *A*(*ρ_dg_,ρ_cv_*, 0) is Hurwitz stable since we have that *A_ρ_* ≤ *A*(*ρ_u_*) and, therefore *λ_PF_* (*A_ρ_*) ≤ *λ_PF_* (*A*(*ρ_u_*)) < 0. Using now the determinant formula, we have that

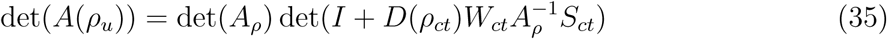

where *D*(*ρ_ct_*) := diag(*ρ_ct_*) and *W_ct_* is defined such that diag(*ρ_ct_*)*W_ct_x* is the vector of propensity functions associated with the catalytic reactions. Hence, this implies that

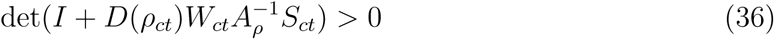

for all 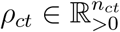. Since the matrices *W_ct_,S_ct_* are nonnegative, the diagonal entries of *D*(*ρ_ct_*) are positive and 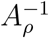 is nonpositive (since *A_ρ_* is Metzler and Hurwitz stable), then it is necessary that all the eigenvalues of 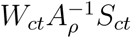 be zero for the determinant to remain positive. This completes the argument.

The converse (i.e. (d) implies (c)) can be proven by noticing that if *A_ρ_* is Hurwitz stable, then *A_ρ_* + *ϵS_ct_W_ct_* remains Hurwitz stable for some sufficiently small *ϵ* > 0. This proves the existence of a 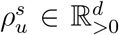 such that the matrix 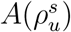. Using the determinant formula, it is immediate to see that the second statement implies the determinant condition of statement (c).

The equivalence between the statements (d) and (e) comes from Lemma 20 and the fact that the sign-pattern of the inverse of a Hurwitz stable Metzler matrix is uniquely defined by its sign-pattern.

Let us now focus on the equivalence between the statements (d) and (f) under the assumption that *S_cv_* contains exactly one entry equal to −1 and one equal to 1. Assume w.l.o.g that 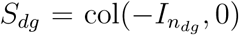. Then, we have that 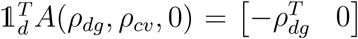. Hence, the function 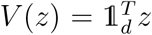 is a weak Lyapunov function for the linear positive system 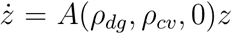. Invoking LaSalle’s invariance principle, we get that the matrix is Hurwitz stable if and only if the matrix

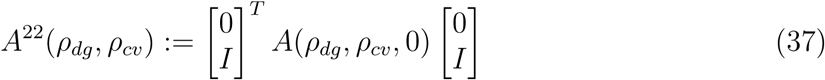

is Hurwitz stable for all 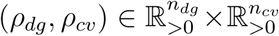. Note that this is a necessary condition for the matrix *A*(*ρ_dg_,ρ_cv_*, 0) to be Hurwitz stable for all rate parameters values. Hence, this means that the stability of the matrix *A_ρ_* is equivalent to the Hurwitz stability of 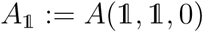. Finally, since *A*^22^(*ρ_dg_,ρ_cv_*) is Hurwitz stable, then we have that 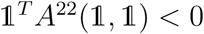. The proof is complete.

## 7 Examples

### 7.1 Example 1: Modified stochastic switch

We propose to illustrate the results by considering a variation of the stochastic switch [51] described by the set of reactions given in Table 1, where the functions *f*_1_ and *f*_2_ are valid nonnegative functions (e.g. mass-action or Hill-type). Our goal is to control the mean population of ***X*_2_** by actuating ***X***_1_. We further assume that *α*_1_*,α*_2_*,γ*_1_*,γ*_2_ > 0, which implies that the state-space of the open-loop network is irreducible.

#### Scenario 1.

In this scenario, we simply assume that *f*_1_ and *f*_2_ are bounded functions with respective upper-bounds *β*_1_ > 0 and *β*_2_ > 0. Then, using the results in [15], the ergodicity of the network in Table 1 can be established by checking the ergodicity of a comparison network obtained by substituting the functions *f*_1_ and *f*_2_ by their upper-bound. In the current case, the comparison network coincides with a unimolecular network with mass-action kinetics defined with

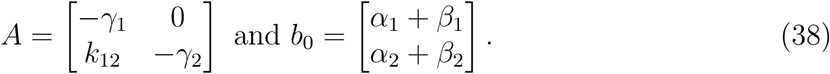

**Table 1:** Modified stochastic switch of [51]

|  | Reaction | Propensity |
| --- | --- | --- |
| $\mathcal{R}_1$ | $\emptyset \longrightarrow \mathbf{X}_1$ | $\alpha_1 + f_1(X_2)$ |
| $\mathcal{R}_2$ | $\mathbf{X}_1 \longrightarrow \emptyset$ | $\gamma_1 X_1$ |
| $\mathcal{R}_3$ | $\emptyset \longrightarrow \mathbf{X}_2$ | $\alpha_2 + f_2(X_1) + k_{12} X_1$ |
| $\mathcal{R}_4$ | $\mathbf{X}_2 \longrightarrow \emptyset$ | $\gamma_2 X_2$ |

It is immediate to see that the characteristic matrix is Hurwitz stable and that the system is output controllable provided that *k*_12_ ≠ 0 since 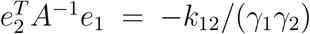 (see Lemma 6). Hence, tracking is achieved provided that the lower bound condition (9) in Theorem 7 is satisfied. Moreover, we can see that for any *α*_1_*,α*_2_*,β*_1_*,β*_2_*,γ*_1_*,γ*_2_*,k*_12_ > 0, we have that 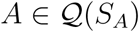 and 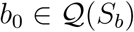 where

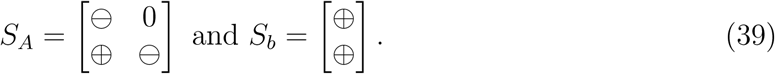

All the matrices in 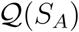 are Hurwitz stable since the matrix sgn(*S_A_*) is Hurwitz stable or, equivalently, since the graph associated with *S_A_* given by 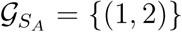 is acyclic (Lemma 17). Moreover, this graph trivially contains a path from node 1 to node 2, proving then that tracking will be ensured by the AIC motif provided that the inequality (32) is satisfied (Theorem 18). Alternatively, we can prove this by augmenting the graph 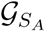 with the edge {(2, 1)} (see the proof of Theorem 18) to obtain the graph 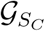 with associated sign matrix

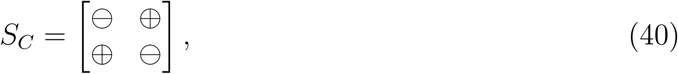

which is not sign-stable since the matrix sgn(*S*C) has an eigenvalue located at 0 or, equivalently, since the graph 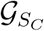 has a cycle (Lemma 17). To arrive to the same result, we can also check that the vectors *v*_1_ = (2, 1), *v*_2_ = (0, 0) and *v*_3_ = (1/2, 1/2) solve the linear program (33).

#### Scenario 2.

We slightly modify here the previous scenario by making the function *f*_1_ affine in *X*_2_, i.e. *f*_1_(*X*_2_)= *k*_21_*X*_2_ + *δ*_1_, *k*_21_*,δ*_1_ ≥ 0. It is immediate to see that the structural result fails as the resulting characteristic matrix has the same sign pattern as the matrix *S_C_*. Hence, the network is not structurally ergodic but it can be robustly ergodic. To illustrate this, we define the following intervals for the parameters *γ*_1_ ∈ [1, 2], *γ*_2_ ∈ [3, 4], 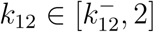 and 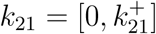, where 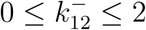 and 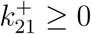. Hence, we have that

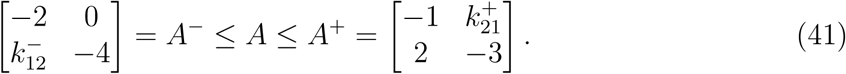

The stability of all the matrices *A* in this interval is established through the stability of *A*^+^ (Lemma 8), which holds provided that 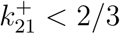. This can also be checked by verifying that *v^T^ A* < 0 for 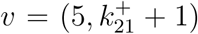 under the very same condition on 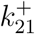. Regarding the output controllability, we need to consider the matrix *A*^−^ (Theorem 9) and observe that if 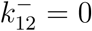 then output controllability does not hold as there is no path from node 1 to node 2 in the graph (Lemma 6). Alternatively, we can check that, in this case, 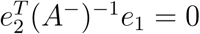 (Theorem 9) or that the linear program (18) is not feasible. To conclude, when 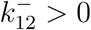 and 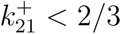 then the linear program (18) is feasible with the vectors 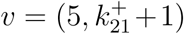, and 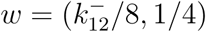, proving then that, in this case, the AIC motif will ensure robust tracking for the controlled network provided that the condition (14) is satisfied.

### 7.2 Example 2: SIR model

Let us consider the open stochastic SIR model considered in [15] described by the matrices

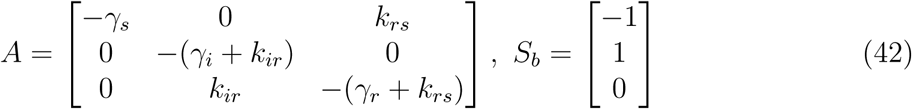

where all the parameters are positive. The constraint *v^T^ S_b_* = 0 enforces that 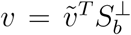, 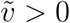, where 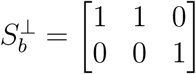. This leads to

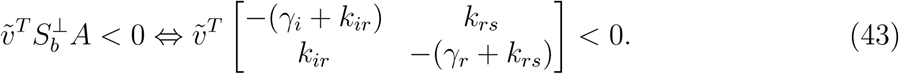

Since the entries are not independent, the use of sign-matrices or interval matrices are conservative. However, if we use Theorem 21, then we can just substitute the parameters by 1 and observe that the resulting matrix is Hurwitz stable to prove the structural stability of the matrix. Alternatively, we can take 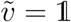 and obtain

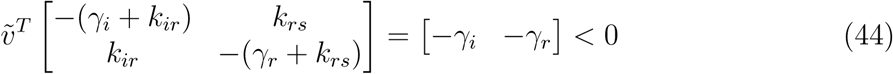

from which the same result follows.

### 7.3 Example 3: Circadian Clock

We consider the circadian clock-model of [10] which is described by the matrices

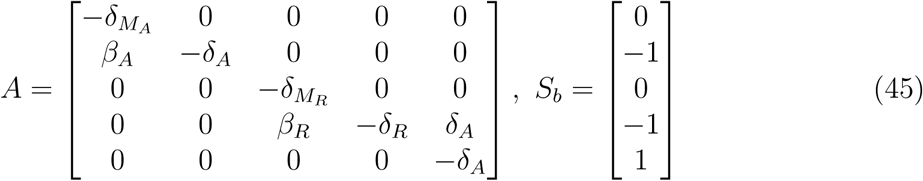

where all the parameters are positive. As in the previous example, the condition reduces to

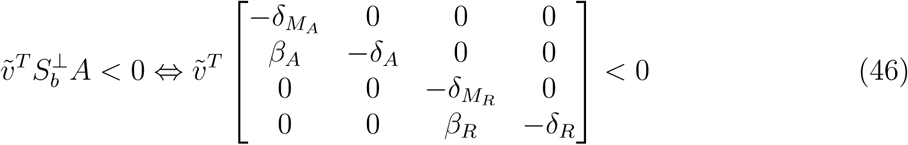

where 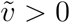. Clearly, we have four degradation reactions and two catalytic ones. Using the last statement of Theorem 21 we get that the matrix 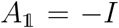. We also have in this case that

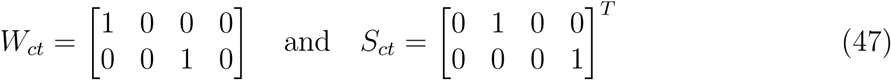

and, hence, 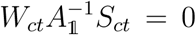. Hence, the system is structurally stable. Alternatively, the triangular structure of the matrix would also lead to the same conclusion.

### 7.4 Example 4: Toy model

Let us consider here the following toy network where

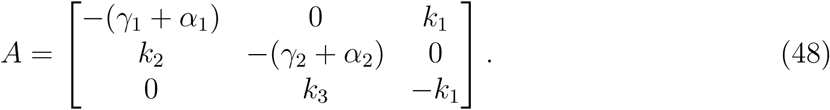

Assume that *α*_1_ = *k*_2_ and *α*_2_ = *k*_3_. Then, we get that

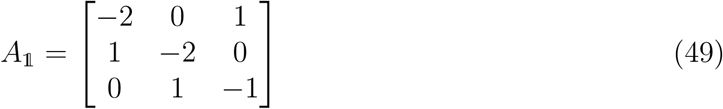

is Hurwitz stable and hence that the matrix is structurally stable. However, if we assume now that *α*_1_ = *α*_2_ = 0, then we get

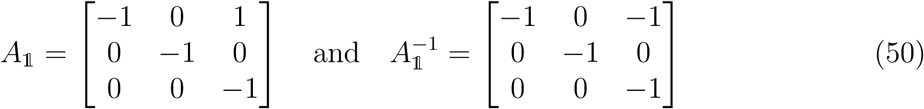

where 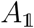 is Hurwitz stable. We have in this case that

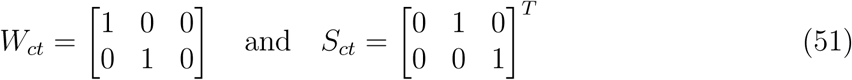

and hence

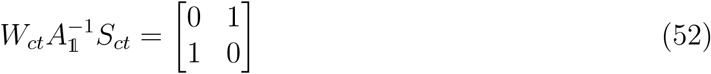

which has a spectral radius equal to 1. Hence, the matrix is not structurally stable. Define now

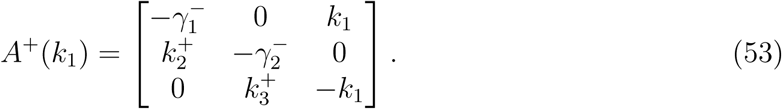

Using a perturbation argument, we can prove that the 0-eigenvalue of *A*^+^(0) locally bifurcates to the open right-half plane for some sufficiently small *k*_1_ > 0 if and only if *k*_2_*k*_3_ − *γ*_1_*γ*_2_ < 0. Hence, there exists a *k*_1_ > 0 such that *A*^+^(*k*_1_) is Hurwitz stable if and only if *k*_2_*k*_3_ − *γ*_1_*γ*_2_ < 0. Noting now that

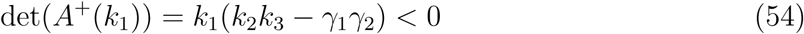

and, hence, the determinant never switches sign, which proves that the matrix *A*^+^(*k*_1_) is structurally stable.

## 8 Conclusion

Several extensions of nominal ergodicity and output controllability results initially proposed in [3, 15] have been obtained. Those extensions are based on the use of interval matrices, sign-matrices and parameter-dependent matrices, respectively. The two first approaches have the benefits of leading to conditions that are easy to check and applicable to large networks as the complexity of the conditions scales linearly with the number of species. However, they may suffer from an arbitrarily large conservatism as the interval matrix and the sign-matrix representation is not necessarily exact. This issue is palliated by the use of parameter-dependent matrix where the entries of the characteristic matrix are exactly parameterized in terms of the reaction rates. This representation is exact but is more difficult to theoretically exploit. Numerically, it is known that checking the stability of a parameter-dependent matrix is NP-Hard. However, by exploiting the Metzler structure of the matrix, it has been possible to obtain interesting simplified conditions for the robust and structural ergodicity of stochastic reaction networks with uncertain reaction rates. Possible extensions of those results would pertain on more general reaction networks or the development of efficient algorithms for the analysis and the control of reaction networks.

## Footnotes

1 A Metzler matrix is a square matrix with nonnegative off-diagonal elements.

2 Computationally tractable conditions for checking the irreducibility of reaction networks are provided in [41].

## References

[1] C. Briat and M. Khammash, “Robust ergodicity and tracking in antithetic integral control of stochastic biochemical reaction networks,” in 55th IEEE Conference on Decision and Control, Las Vegas, USA, 2016, pp. 752–757.

[2] C. Briat and M. Khammash, “Robust and structural ergodicity analysis of stochastic biomolecular networks involving synthetic antithetic integral controllers,” in 20th IFAC World Congress, Toulouse, France, 2017, pp. 11 405–11 410.

[3] C. Briat, A. Gupta, and M. Khammash, “Antithetic integral feedback ensures robust perfect adaptation in noisy biomolecular networks,” Cell Systems, vol. 2, pp. 17–28, 2016.

[4] D. K. Ro, E. M. Paradise, M. Ouellet, K. J. Fisher, K. L. Newman, J. M. Ndungu, K. A. Ho, R. A. Eachus, T. S. Ham, J. Kirby, M. C. Y. Chang, S. T. Withers, Y. S. R. Sarpong, and J. D. Keasling, “Production of the antimalarial drug precursor artemisinic acid in engineered yeast,” Nature, vol. 440(7086), pp. 940–943, 2006.

[5] M. Feinberg, “Complex balancing in general kinetic systems,” Archive for rational mechanics and analysis, vol. 49(3), pp. 187–194, 1972.

[6] F. Horn and R. Jackson, “General mass action kinetics,” Archive for rational mechanics and analysis, vol. 47(2), pp. 81–116, 1972.

[7] J. Goutsias and G. Jenkinson, “Markovian dynamics on complex reaction networks,” Physics reports, vol. 529, pp. 199–264, 2013.

[8] M. Thattai and A. van Oudenaarden, “Intrinsic noise in gene regulatory networks,” Proceedings of the National Academy of Sciences, vol. 98(15), pp. 8614–8619, 2001.

[9] M. B. Elowitz, A. J. Levine, E. D. Siggia, and P. S. Swain, “Stochastic gene expression in a single cell,” Science, vol. 297(5584), pp. 1183–1186, 2002.

[10] J. M. G. Vilar, H. Y. Kueh, N. Barkai, and S. Leibler, “Mechanisms of noise-resistance in genetic oscillator,” Proc. Natl. Acad. Sci., vol. 99(9), pp. 5988–5992, 2002.

[11] J. Paulsson, O. G. Berg, and M. Ehrenberg, “Stochastic focusing: Fluctuation-enhanced sensitivity of intracellular regulation,” Proceedings of the National Academy of Sciences, vol. 97(13), pp. 7148–7153, 2000.

[12] A. Gupta, B. Hepp, and M. Khammash, “Noise induces the population-level entrainment of incoherent, uncoupled intracellular oscillators,” Cell Systems (to appear), 2016.

[13] D. Anderson and T. G. Kurtz, “Continuous time Markov chain models for chemical reaction networks,” in Design and analysis of biomolecular circuits - Engineering Approaches to Systems and Synthetic Biology, H. Koeppl, D. Densmore, G. Setti, and M. di Bernardo, Eds. Springer Science+Business Media, 2011, pp. 3–42.

[14] D. F. Anderson and T. G. Kurtz, Stochastic Analysis of Biochemical Systems, ser. Mathematical Biosciences Institute Lecture Series. Springer Verlag, 2015, vol. 1.2.

[15] A. Gupta, C. Briat, and M. Khammash, “A scalable computational framework for establishing long-term behavior of stochastic reaction networks,” PLOS Computational Biology, vol. 10(6), p. e1003669, 2014.

[16] D. Del Vecchio, A. J. Dy, and Y. Qian, “Control theory meets synthetic biology,” Journal of The Royal Society Interface, vol. 13, no. 120, 2016.

[17] G. Lillacci, S. K. Aoki, D. Schweingruber, and M. Khammash, “A synthetic integral feedback controller for robust tunable regulation in bacteria,” bioRxiv, 2017. [Online]. Available: http://www.biorxiv.org/content/early/2017/08/01/170951

[18] C. Briat, A. Gupta, and M. Khammash, “Variance reduction for antithetic integral control of stochastic reaction networks,” Journal of the Royal Society: Interface, vol. 15(143), p. 20180079, 2018.

[19] C. Briat, A. Gupta, I. Shames, and M. Khammash, “Scalable tests for ergodicity analysis of large-scale interconnected stochastic reaction networks,” in 21st International Symposium on Mathematical Theory of Networks and Systems, Groningen, The Netherlands, 2014, pp. 92–95.

[20] R. E. Moore, R. B. Keafott, and M. J. Cloud, Introduction to Interval Analysis. SIAM, 2009.

[21] C. Jeffries, V. Klee, and P. van den Driessche, “When is a matrix sign stable?” Canadian Journal of Mathematics, vol. 29, pp. 315–326, 1977.

[22] R. A. Brualdi and B. L. Shader, Matrices of sign-solvable linear systems. Cambridge, UK: Cambridge University Press, 1995.

[23] C. Briat, “Sign properties of Metzler matrices with applications,” Linear Algebra and its Applications, vol. 515, pp. 53–86, 2017.

[24] M. McCreesh and B. Gharesifard, “Stability of bounded subsets of metzler sparse matrix cones,” Automatica, vol. 95, pp. 544–547, 2018.

[25] J. W. Helton, I. Klep, and R. Gomez, “Determinant expansions of signed matrices and of certain jacobians,” SIAM Journal on Matrix Analysis and Applications, vol. 31(2), pp. 732–754, 2009.

[26] J. W. Helton, V. Katsnelson, and I. Klep, “Sign patterns for chemical reaction networks,” Journal of Mathematical Chemistry, vol. 47, pp. 403–429, 2010.

[27] C. Briat, “Sign properties of Metzler matrices with applications,” Linear Algebra and its Applications, vol. 515, pp. 53–86, 2017.

[28] G. Giordano, C. Cuba Samaniego, E. Franco, and F. Blanchini, “Computing the structural influence matrix for biological systems,” Journal of Mathematical Biology, vol. 72, pp. 1927–1958, 2016.

[29] F. Horn and R. Jackson, “General mass action kinetics,” Archive for Rational Mechanics and Analysis, vol. 47, pp. 81–116, 1972.

[30] D. F. Anderson and J. Kim, “Some network conditions for positive recurrence of stochastically modeled reaction networks,” SIAM Journal on Applied Mathematics, vol. 78(5), pp. 2692–2713, 2018.

[31] J. C. Doyle, “Analysis of feddback systems with structured uncertainties,” IEEE Proc., Part D, vol. 129, pp. 242–250, 1982.

[32] A. Packard and J. C. Doyle, “The complex structured singular value,” Automatica, vol. 29, pp. 71–109, 1993.

[33] M. H. Khammash, “Necessary and sufficient conditions for the robustness of time-varying systems with applications to sampled-data systems,” IEEE Transactions on Automatic Control, vol. 38(1), pp. 49–57, 1993.

[34] W. Michiels and S. I. Niculescu, Stability and stabilization of time-delay systems. An eigenvalue based approach. Philadelphia, USA: SIAM Publication, 2007.

[35] A. P. Seyranian and A. A. Mailybaev, Multiparameter stability theory with mechanical applications. Singapore: World Scientific, 2003.

[36] C. Briat, Linear Parameter-Varying and Time-Delay Systems – Analysis, Observation, Filtering & Control, ser. Advances on Delays and Dynamics. Heidelberg, Germany: Springer-Verlag, 2015, vol. 3.

[37] F. Blanchini and S. Miani, “A New Class of Universal Lyapunov Functions for the Control of Uncertain Linear Systems,” IEEE Transactions on Automatic Control, vol. 44(3), pp. 641–647, 1999.

[38] D. Angeli, P. De Leenheer, and E. D. Sontag, “Chemical networks with inflows and outflows: A positive linear differential inclusions approach,” Biotechnology Progress, vol. 25(3), pp. 632–642, 2009.

[39] F. Blanchini and G. Giordano, “Piecewise-linear Lyapunov functions for structural stability of biochemical networks,” Automatica, vol. 50(10), pp. 2482–2493, 2014.

[40] G. Giordano, “Structural analysis and control of dynamical networks,” Ph.D. dissertation, University of Udine, 2016.

[41] A. Gupta and M. Khammash, “Determining the long-term behavior of cell populations: A new procedure for detecting ergodicity in large stochastic reaction networks,” ETH-Zürich, Tech. Rep. arXiv:1312.2879, 2013.

[42] A. Berman and R. J. Plemmons, Nonnegative matrices in the mathematical sciences. Philadelphia, USA: SIAM, 1994.

[43] W. Mitkowski, “Remarks on stability of positive linear systems,” Control and Cybernetics, vol. 29(1), pp. 295–304, 2000.

[44] D. Handelman, “Representing polynomials by positive linear functions on compact convex polyhedra,” Pacific Journal of Mathematics, vol. 132(1), pp. 35–62, 1988.

[45] C. Briat, “Robust stability and stabilization of uncertain linear positive systems via integral linear constraints -*L*_1_-and L_∞_-gains characterizations,” International Journal of Robust and Nonlinear Control, vol. 23(17), pp. 1932–1954, 2013.

[46] M. Putinar, “Positive polynomials on compact semi-algebraic sets,” Indiana Univ. Math. J., vol. 42, no. 3, pp. 969–984, 1993.

[47] P. Parrilo, “Structured semidefinite programs and semialgebraic geometry methods in robustness and optimization,” Ph.D. dissertation, California Institute of Technology, Pasadena, California, 2000.

[48] E. Fornasini and M. E. Valcher, “Linear copositive Lyapunov functions for continuous-time positive switched systems,” IEEE Transactions on Automatic Control, vol. 55(8), pp. 1933–1937, 2010.

[49] O. Mason and R. N. Shorten, “On linear copositive Lyapunov functions and the stability of switched positive linear systems,” IEEE Transactions on Automatic Control, vol. 52(7), pp. 1346–1349, 2007.

[50] S. Boyd and L. Vandenberghe, Convex Optimization. Cambridge, MA, USA: Cambridge University Press, 2004.

[51] T. Tian and K. Burrage, “Stochastic models for regulatory networks of the genetic toggle switch,” Proc. Natl. Acad. Sci., vol. 103(22), pp. 8372–8377, 2006.

